# Alignment of time-course single-cell RNA-seq data with CAPITAL

**DOI:** 10.1101/859751

**Authors:** Reiichi Sugihara, Yuki Kato, Tomoya Mori, Yukio Kawahara

**Author notes:** Correspondence should be addressed to Y. Kato.

## Abstract

Recent techniques on single-cell RNA sequencing have boosted transcriptome-wide observation of gene expression dynamics of time-course data at a single-cell scale. Typical examples of such analysis include inference of a pseudotime cell trajectory, and comparison of pseudotime trajectories between different experimental conditions will tell us how feature genes regulate a dynamic cellular process. Existing methods for comparing pseudotime trajectories, however, force users to select trajectories to be compared because they can deal only with simple linear trajectories, leading to the possibility of making a biased interpretation. Here we present CAPITAL, a method for comparing pseudotime trajectories with tree alignment whereby trajectories including branching can be compared without any knowledge of paths to be compared. Computational tests on time-series public data indicate that CAPITAL can align non-linear pseudotime trajectories and reveal gene expression dynamics.

## 1 Introduction

Single-cell RNA-sequencing (scRNA-seq) has enabled us to scrutinize gene expression of dynamic cellular processes such as differentiation, reprogramming and cell death. Since tracing a gene expression level of a certain cell over some period of time is infeasible, *pseudotime* analysis with time course data on a population from a common tissue or an organ is of great value to obtain an approximate landscape of gene expression dynamics in those biological systems.

To model dynamic developmental processes when time-course scRNA-seq data are given, various computational tools were developed to predict a *pseudotime trajectory* [1–5]. The predictable topology of trajectories depends on the method of choice, ranging from a simple linear structure to a tree with branching, and even a complex cycle [6]. A comprehensive comparison among existing trajectory inference tools was also provided to make an exhaustive investigation into their performances [7].

Comparison of pseudotime trajectories will provide a key to unveiling regulators that determine cell fates. For instance, when time-series scRNA-seq data from a disease model are compared with those from a positive control, it is possible to pin down the time point at which the disease begins to manifest itself, thereby discovering differentially expressed genes between the two. Another application of trajectory comparison is to investigate differences in gene expression dynamics between species (e.g. human vs. mouse), which will unravel evolutionary differences in determination of cell fates such as regulation timing for a common gene.

Recently, computational methods for aligning pseudotime trajectories across different experimental conditions have been proposed [8, 9]. They aim to align two linear trajectories with dynamic time warping [10, 11], which is an analogy to the classical problem of pairwise sequence alignment, but differs in that multiple cells in one condition may be matched with one cell in the counterpart at a time point. It remains, however, an unsettled question how one can select trajectories to be compared if the pseudotime trajectories include branching, as the above methods based on dynamic time warping can deal only with linear trajectories for comparison. Given that a fair number of cellular processes would include branching, there is still a possibility of making a wrong biological interpretation in a biased manner.

To tackle this problem, dynamic time warping on a pair of linear trajectories has been extended so that it can find the best alignment between all possible pairs of paths with the common progenitor-descendant relationship in the trajectory trees [12]. More precisely, an arboreal matching between the two trajectory trees, which allows unmatched nodes, is computed in the framework of integer linear programing with a branch-and-cut strategy. This approach requires linear programming relaxation for practical use on a large number of single cells due to the wide expressive power of the original problem formulation, which means that approximated results have to be carefully investigated.

Another approach has been proposed to integrate different data sets and estimate cell differences in gene expression across data using a variant of autoencoder [13]. Although this method can cope with multiple scRNA-seq data generated with different conditions, it is designed to interpolate the expression profile of cells to reveal the difference of the same cell state across conditions. This means that the main purpose of their study is not to consider the difference between dynamic cell states along trajectories but to embeds an admixture of cells from different conditions onto the same low dimensional space. In addition, it needs elaborate modeling of a network structure and training of a number of hyper parameters.

In this article, we present a computational method for <u>c</u>omparative <u>a</u>nalysis of <u>p</u>seudotime trajectory inference with tree <u>al</u>ignment (CAPITAL). When a pair of time-series scRNA-seq data sets generated with different experimental conditions is given, CAPITAL seeks to infer respective pseudotime trajectories that may include branching, and then to compute an alignment between the two trajectories. This means that CAPITAL can cope with tree shapes for alignment without selecting trajectories to be compared. Of note, a tree alignment algorithm that CAPITAL employs can output an exact optimal solution rather than an approximate one as in the literature [12]. Computational tests on time-course scRNA-seq data indicate that CAPITAL can align potential tree trajectories as well as viewing gene expression dynamics along pseudotime, which could provide a key to unravelling novel regulators that determine cell fates.

## 2 Methods

### 2.1 Overview of CAPITAL

Taking a pair of expression profiles from two experimental conditions as input, CAPITAL aims to compute respective pseudotime trajectories and align them even if the trajectories include branching (Fig. 1a). Of note, the method deals with clusters rather than constituent cells during alignment to minimize the effect of an inherent noise from scRNA-seq data. After any aligned paths from the resulting alignment are chosen, cells derived from the corresponding clusters are ordered by pseudotime (Fig. 1b). For a specific gene or a set of genes, the difference in expression dynamics between two conditions can be investigated via dynamic time warping.

**Fig. 1.**
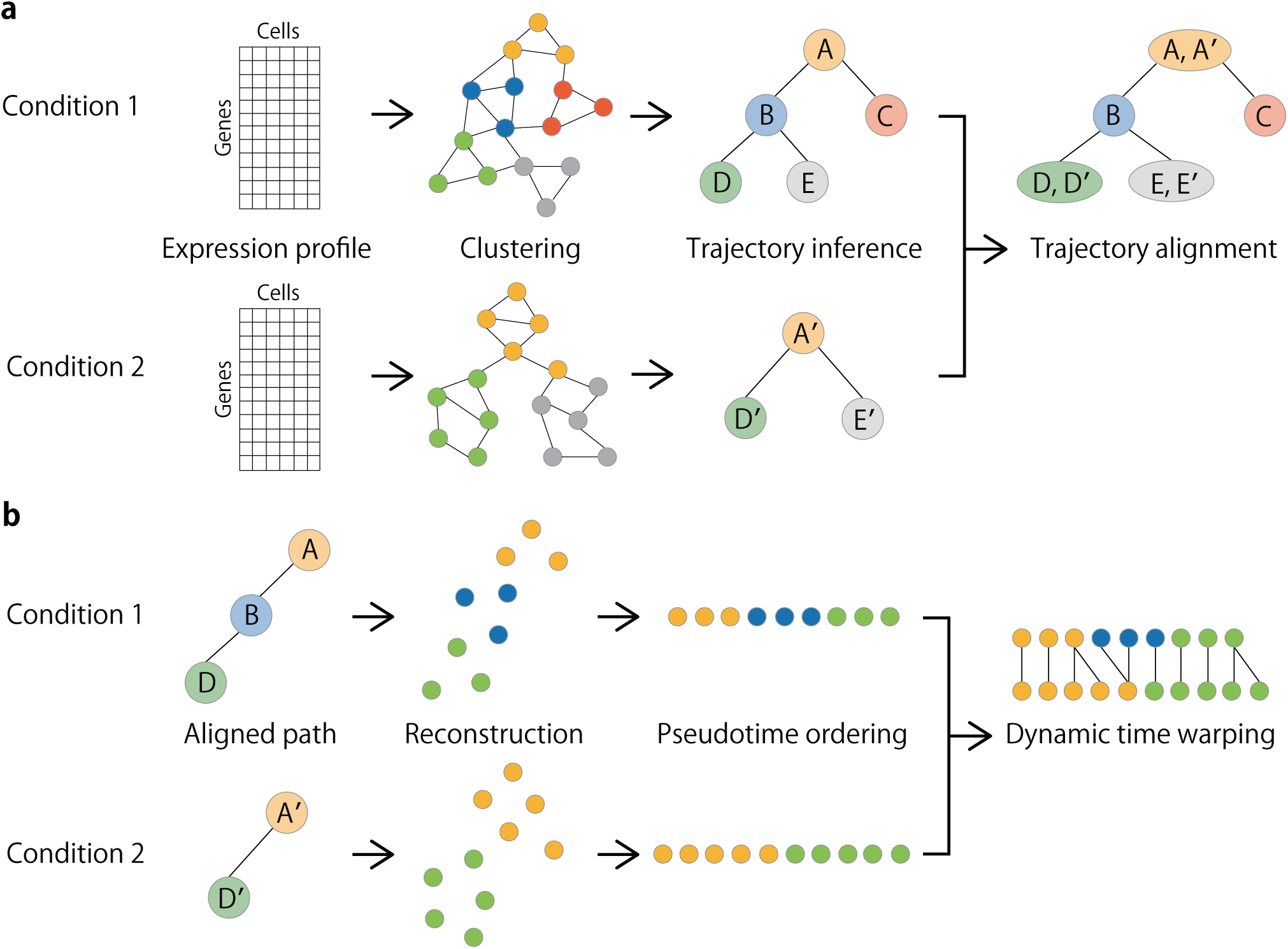
Overview of CAPITAL. **a**, From an expression profile in each experimental condition, a nearest neighbor graph representing cells and their similarities in the expression space as vertices and weighted edges, respectively, is constructed, and then clustered via community detection. Next, the clustered graph is converted into a PAGA graph with clusters and their connectivities, along with cluster centroids, which is used to infer a trajectory by computing a minimum spanning tree. Finally, a trajectory alignment is obtained by aligning one minimum spanning tree with the other across different conditions. **b**, When a pair of aligned paths from the trajectory alignment is taken, clusters in each condition are decomposed into their original cells, and the start cell in the first cluster (e.g. A and A’) on the aligned path is determined in a way that it has the longest distance to a cell in another cluster. Computing the accumulated transition matrix for these cells with the start cell generates a pseudotime order. Dynamic time warping is then performed for a gene of interest to investigate the dynamic relationship among cells for that gene along the pseudotime.

### 2.2 Clustering cells via graph structure

Let *X* = (***x***_1_, …, ***x***_*n*_) ∈ ℝ^*m*×*n*^ be an expression profile matrix of *m* genes and *n* cells, where ***x***_*j*_ = (*x*_1*j*_, …, *x*_*mj*_)^T^ ∈ ℝ^*m*^ (1 ≤ *j* ≤ *n*) is the logarithm of a normalized count vector of cell *j* with respect to genes 1, …, *m*. Note that the genes may be assumed to be highly variable in expression over all cells. A weighted graph is constructed in a way that each vertex represents a cell, and each edge connecting two cells *i* and *j* has a weight calculated by the Euclidian distance ‖***x***_*i*_−***x***_*j*_‖ between the two cells. From this graph, a *k*-nearest neighbor (*k*-NN) graph can be built on the basis of the distance assigned to each edge, as it can better capture phenotypic relatedness [14]. Clustering vertices in the *k*-NN graph is then performed by a community detection method such as the Leiden algorithm [15].

### 2.3 Inferring a pseudotime trajectory

To make the graph structure simple, a partition-based graph abstraction (PAGA) graph [5] is constructed from the clustered *k*-NN graph, where each cluster represents a vertex and each edge connecting two vertices has an associated connectivity. In addition, a cluster centroid is defined as the cell whose distance to the other cells in the same cluster is minimum. This is useful for aligning clusters across conditions since each cluster in the PAGA graph is a conceptual object and does not have an actual expression vector. Finally, a minimum spanning tree in each PAGA graph is computed with Kruskal’s algorithm. If one chooses one of the centroids as the root of the tree, the resulting rooted minimum spanning tree can be regarded as a pseudotime trajectory tree. Of note, actual pseudotime of each cell along a path in the tree will be estimated later.

### 2.4 Preliminaries to handling trees

Let *T* = (*V, E*) be an unordered labeled rooted tree, where *V* and *E* (also denoted by *V* (*T*) and *E*(*T*)) are a set of nodes (vertices) and that of edges of the tree, respectively, and the children of each node are regarded as a set. The number of nodes in tree *T* is represented as |*T*|. Let *θ* be the empty tree, and let *T* (*i*) be the subtree of *T* rooted at node *i* ∈ *V* (*T*). If node *i* has children *i*_1_, …, *i*_*m*_, i.e., the degree of node *i* is *m*, we define *F* (*i*) = *F* (*i*_1_, …, *i*_*m*_) as the forest comprising subtrees *T* (*i*_1_), …, *T* (*i*_*m*_).

Inserting node *w* ∈ *V* (*T*) as a child of node *v* ∈ *V* (*T*) makes *w* be the parent of a consecutive subset of the children of node *v*. An alignment of unordered labeled trees *T*_1_ and *T*_2_ is obtained by inserting nodes labeled with *spaces λ* into *T*_1_ and/or *T*_2_ so that the two trees are isomorphic if the labels are ignored, and then by overlaying the resulting trees. Let *γ* : (Σ_*λ*_, Σ_*λ*_) \ ({*λ*}, {*λ*}) → ℝ denote a metric cost function on pairs of labels, where Σ is a finite alphabet and Σ_*λ*_ = Σ ∪ {*λ*}. We often extend this notation to nodes where *γ*(*i, j*) means *γ* (label(*i*), label(*j*)) for *i, j* ∈ *V* (*T*). For single-cell analysis, we define the cost function as

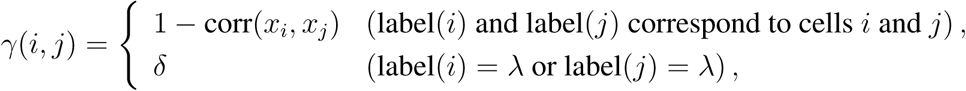

where corr(*x*_*i*_, *x*_*j*_) returns Spearman’s rank correlation coefficient between expression levels *x*_*i*_ and *x*_*j*_ of cells *i* and *j*, respectively, and *δ* is some constant.

The cost of the alignment is defined as the sum of the costs of all paired labels in the alignment. An *optimal alignment* of *T*_1_ and *T*_2_ is an alignment with the minimum cost, which is called the optimal *alignment distance* between *T*_1_ and *T*_2_ denoted by *D*(*T*_1_, *T*_2_). This notion is extended to two forests *F*_1_ and *F*_2_, and denoted by *D*(*F*_1_, *F*_2_).

### 2.5 Tree alignment algorithm

The problem of computing an optimal alignment distance between general unordered trees is MAX SNP-hard [16]. However, aligning unordered trees with ‘bounded’ degrees can be solved in polynomial time. In what follows, we will see the dynamic programming (DP) algorithm for calculating the optimal alignment distance between unordered trees with bounded degrees presented in [16].

#### 2.5.1 Initialization

The initial settings for handling the empty tree during the DP calculation are defined as follows:

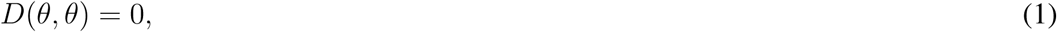

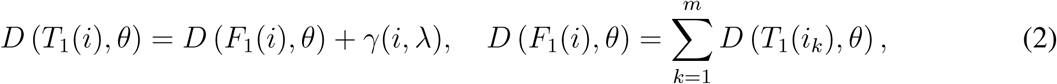

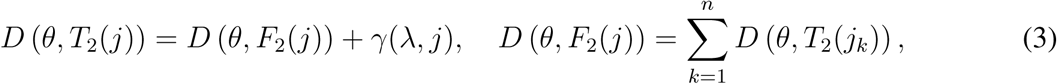

where *m* and *n* are degrees of nodes *i* ∈ *V* (*T*_1_) and *j* ∈ *V* (*T*_2_), respectively.

#### 2.5.2 Recursion

We first look at the computation of an optimal alignment distance between trees.

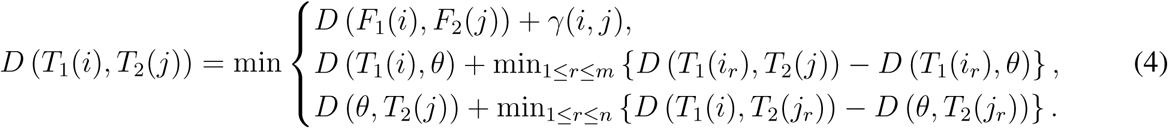

This recursion means that there are three cases to be considered for the DP calculation: (i) paired nodes (*i, j*) is in the alignment; (ii) (*i, λ*) and (*k, j*) for some *k* ∈ *V* (*T*_1_) are in the alignment; and (iii) (*λ, j*) and (*i, h*) for some *h* ∈ *V* (*T*_2_) are in the alignment. A proof of the correctness of this recursion can be found in [16, 17].

We next focus on how to compute an optimal alignment distance *D*(*F*_1_(*i*), *F*_2_(*j*)) between forests appeared in Equation (4). Since all combinations of forest pairs derived from unordered trees *T*_1_(*i*) and *T*_2_(*j*) are needed to be considered, we define subsets of forests denoted by 𝒜 ⊆ {*T*_1_(*i*_1_), …, *T*_1_(*i*_*m*_)}and ℬ ⊆ {*T*_2_(*j*_1_), …, *T*_2_(*j*_*n*_)}. The alignment distance between forest sets 𝒜 and ℬ can then be computed by 

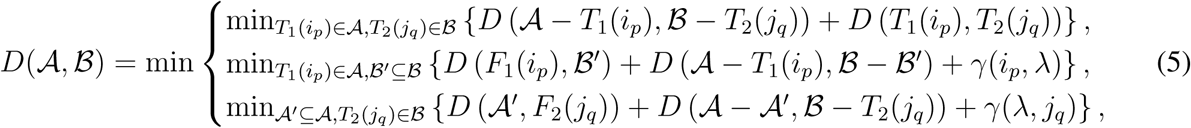

where 1 ≤ *p* ≤ *m* and 1 ≤ *q* ≤ *n*. Note that *D*(𝒜, ℬ) where 𝒜 = {*T*_1_(*i*_1_), …, *T*_1_(*i*_*m*_)} and ℬ = {*T*_2_(*j*_1_), …, *T*_2_(*j*_*n*_)} is equivalent to *D*(*F*_1_(*i*), *F*_2_(*j*)). Since degrees *m* and *n* are bounded and regarded as constants in this case, time and space complexities of the tree alignment algorithm are evaluated as *O*(|*T*_1_||*T*_2_|).

#### 2.5.3 Pseudocode

Assume that the nodes in tree *T*_*k*_ (*k* = 1, 2) are numbered by 1 through |*T*_*k*_| in the postorder fashion. The following pseudocode computes an optimal tree alignment distance between the ordered trees with bounded degrees:

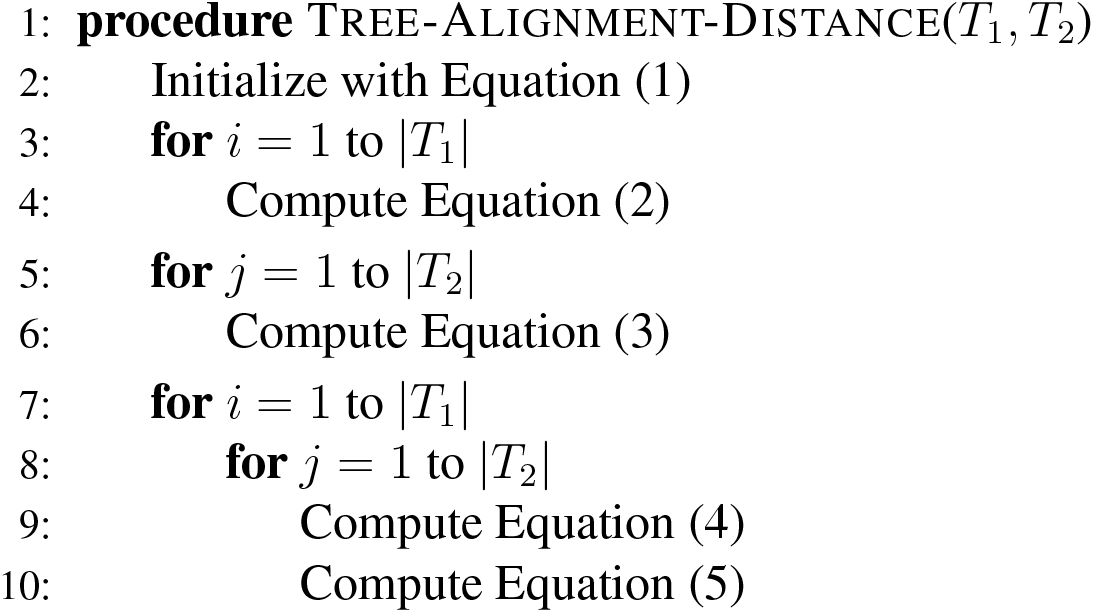

#### 2.5.4 Traceback

*D*(*T*_1_(|*T*_1_|), *T*_2_(|*T*_2_|)) obtained in the above pseudocode will have an optimal alignment distance between *T*_1_ and *T*_2_. To recover the optimal tree alignment, the traceback procedure starting with *D*(*T*_1_(|*T*_1_|), *T*_2_(|*T*_2_|)) is needed, where the calculation path to *D*(*T*_1_(|*T*_1_|), *T*_2_(|*T*_2_|)) is recovered with traceback pointers that hold the choice of the minimum operations in Equations (4) and (5).

### 2.6 Pseudotime ordering

Having been obtained from a trajectory alignment between two conditions, a pair of aligned paths is arbitrarily chosen, and all cells in a cluster are to be processed in each condition. Note that the start cell in the first cluster on the aligned path has to be determined for subsequent pseudotime ordering in a way that it has the longest distance to a cell in another cluster in the expression space. A diffusion pseudotime in each condition is then calculated by taking into consideration the accumulated transition matrix for all cells with respect to the path based on a random walk on a nearest neighbor graph [1].

### 2.7 Dynamic time warping

Dynamic time warping is an algorithm for measuring similarity between temporal sequences. Briefly, given two time series, the algorithm aims to find the best matching between the two sequences by stretching or compressing elements of the sequences. For optimization, the distance between two elements across sequences measured on the basis of time warping functions is summed over all elements, which is then to be minimized by dynamic programming. Details of the algorithm can be found in [11].

### 2.8 Implementation

CAPITAL is implemented with Python, making good use of a single-cell analysis toolkit Scanpy [18].

### 2.9 Preprocessing data

Before aligning time-course scRNA-seq data, each may need to be preprocessed due to noise such as outlier cells included in the original data. In what follows, we will describe how to adapt these data to subsequent comparison.

#### 2.9.1 hFib-MyoD

Human fibroblast MyoD-mediated (hFib-MyoD) reprogramming data [9] were obtained from Gene Expression Omnibus (GEO) database (GSE105211). 658 cells were filtered in a way that cells with at most 200 genes expressed were removed according to Scanpy’s instructions, and outliers revealed with uniform manifold approximation and projection (UMAP) [19] were also removed, resulting in 538 cells. Genes were also filtered so that among genes expressed in at least three cells the top 140 highly variable genes were kept. A PAGA graph with Leiden community was generated from a 8-NN graph, which was used to build a minimum spanning tree.

#### 2.9.2 HSMM

Human skeletal muscle myoblast (HSMM) differentiation data [20] were available at GEO (GSE52529). 372 cells were filtered to remove unrelated cells on the basis of instructions in the literature [9], leading to 111 cells. Genes were filtered in the same way as described above, but the top 120 highly variable genes were used. A 7-NN graph was used for subsequent clustering and trajectory inference.

## 3 Results

### 3.1 Alignment of myogenic reprogramming and myoblast differentiation trajectories

To test how CAPITAL can compare two pseudotime trajectories with potential branchings generated from real time-course data, we used two public scRNA-seq data on human fibroblast MyoD-mediated (hFib-MyoD) reprogramming [9] and human skeletal muscle myoblast (HSMM) differentiation [20], which was first used to quantitatively compare these two models with the early trajectory alignment method [9]. In the following, we will refer to these reprogramming and differentiation data as hFib-MyoD and HSMM, respectively.

We first performed clustering of these scRNA-seq data on the basis of community detection in the nearest neighbor graphs in the CAPITAL framework, and illustrated the clustering results in the two dimensional space via UMAP (Fig. 2). For details of the derivation, refer to Methods Sect. 2.9.

**Fig. 2.**
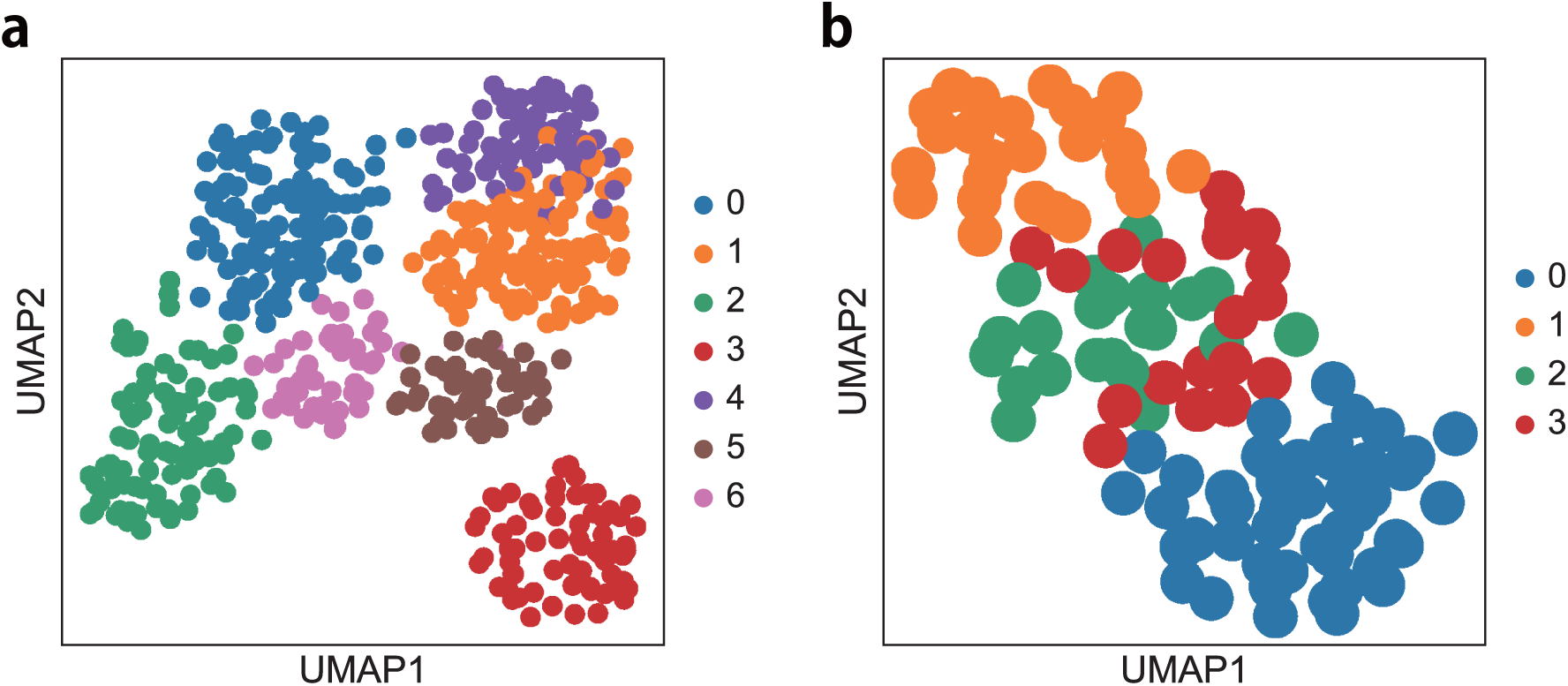
Clustering of scRNA-seq data on myogenic reprogramming and myoblast differentiation embedded in the two dimensional space generated with UMAP. **a**, UMAP for hFib-MyoD. **b**, UMAP for HSMM. Of note, clusters with the same number and color between two experiments do not necessarily mean the same cell type.

Running CAPITAL on the same data, we obtained two minimum spanning trees as predicted pseudotime trajectories in two experiments, which are shown in Fig. 3a,b. A tree alignment between hFib-MyoD and HSMM that CAPITAL computed is illustrated in Fig. 3c, indicating that the algorithm can compute as many matching clusters as possible between different experimental conditions while keeping their pseudotime structures.

**Fig. 3.**
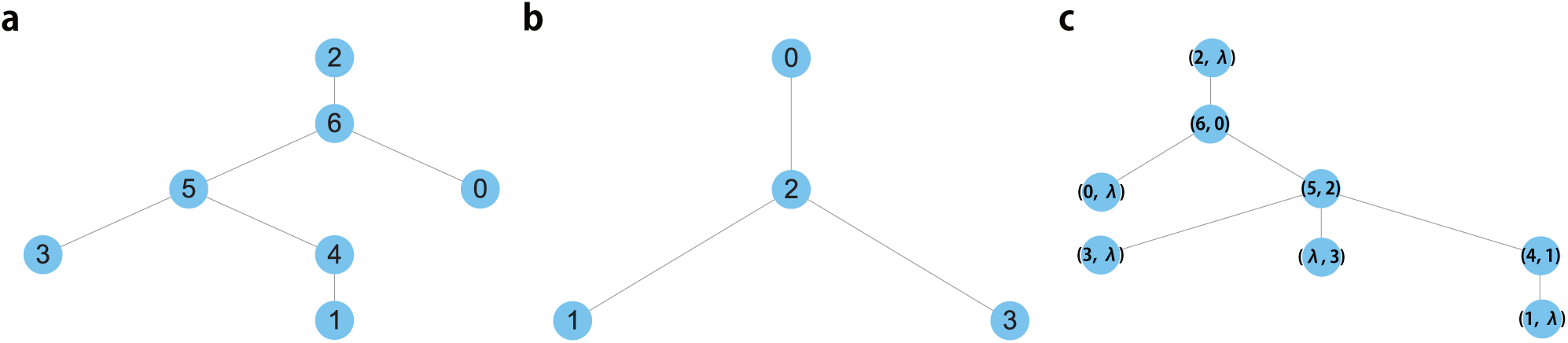
Pseudotime trajectory trees for hFib-MyoD and HSMM computed with CAPITAL. **a**, A trajectory tree for hFib-MyoD. The node drawn on the top of the tree shows the root. **b**, A trajectory tree for HSMM. **c**, A tree alignment between hFib-MyoD and HSMM. Note that a node with a number corresponds to a cluster with the same number in respective data shown in Fig. 2.

Taking a subpath that starts from node (6, 0) and ends with node (4, 1) in the core aligned path from the tree alignment in Fig. 3c, we reconstructed pseudotime orderings for cells included in the subpath within each time-course data with diffusion pseudotime, and computed the matchings between single cells for specific genes along the orderings with dynamic time warping (Fig. 4). Here we chose known marker genes CDK1 for active proliferation, and MEF2C and MYOG for skeletal muscle transcription. As shown in the figure, the relative temporal lag in pseudotime for MYOG tends to be different from those for CDK1 and MEF2C.

**Fig. 4.**
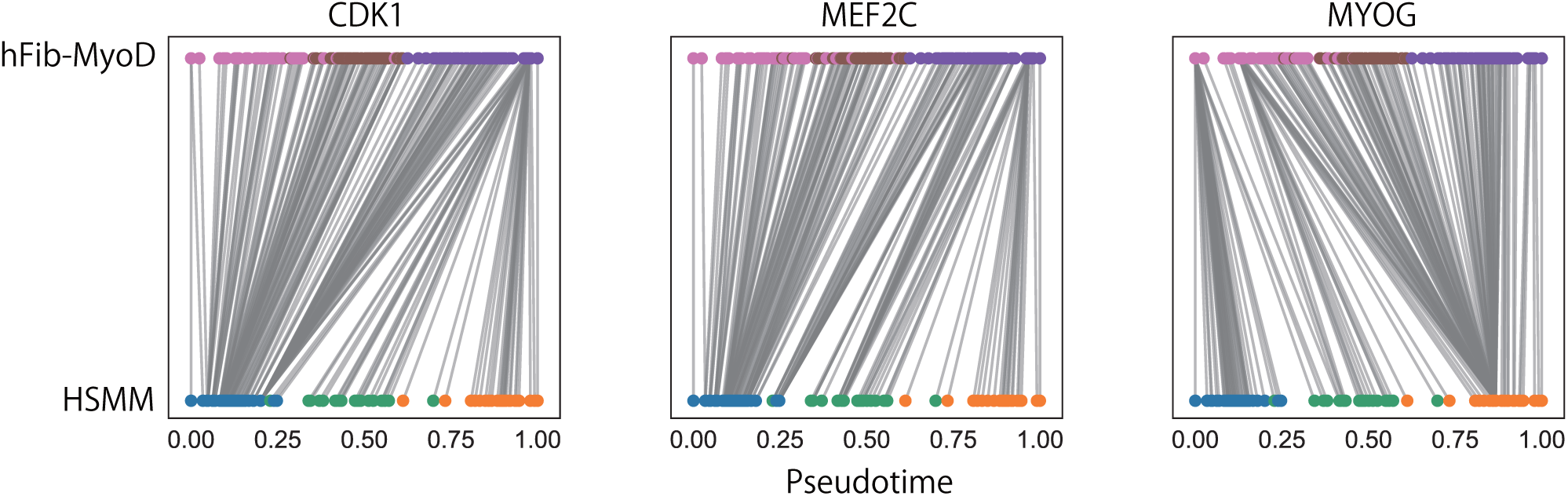
Local matchings between single cells for specific genes along the core aligned path with respect to hFib-MyoD and HSMM. Of note, the coloring for cells in each experiment is the same as in Fig. 2.

Furthermore, we investigated pseudotime kinetics for those marker genes (Fig. 5), which shows a similar tendency to previous results in the literature [9]. These results tell us that CAPITAL can successfully catch the core trajectories to be compared without manually selecting them.

**Fig. 5.**
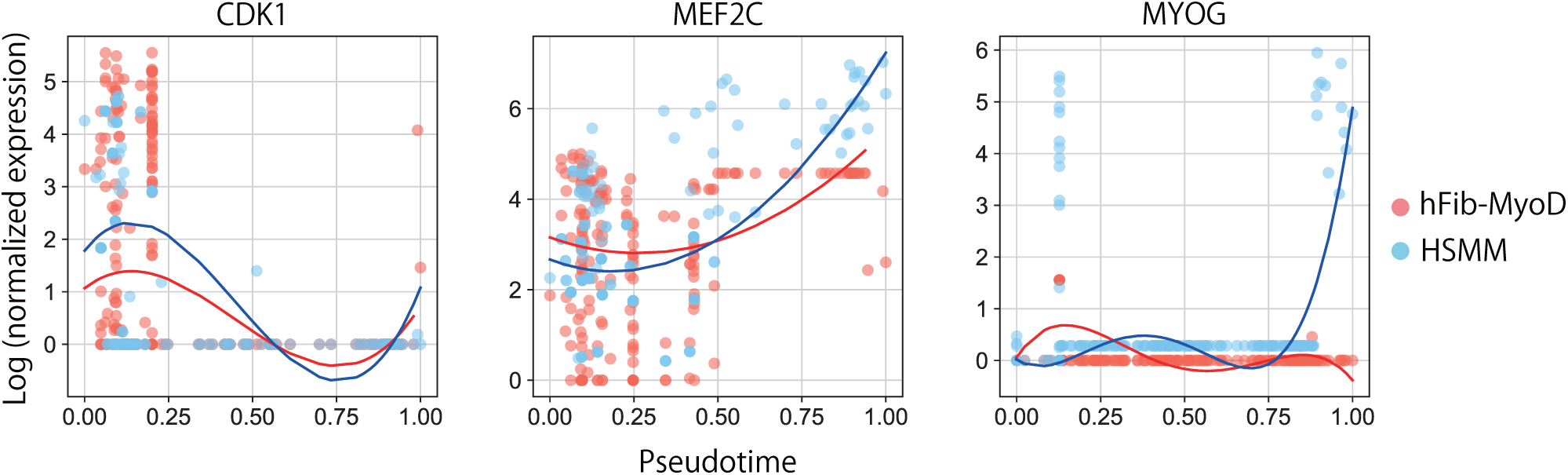
Pseudotime kinetics for marker genes from cells associated with the local matching on the tree alignment between hFib-MyoD and HSMM. Curve fitting with polynomial regression was used to catch a tendency of data points.

## 4 Conclusion

CAPITAL is a computational method for aligning pseudotime trajectories without prior knowledge of trajectory paths to be compared. Hence, there is a possibility of finding out not only regulatory genes on core trajectories but on minor paths. The core part of CAPITAL is to align pseudotime trajectory trees. In this sense, any method for trajectory inference can be incorporated into CAPITAL, which means that it can employ a state-of-the-art or even a future tool with the best prediction accuracy if necessary. Given that scRNA-seq techniques have prevailed in current research in life science, and increasing attention has been paid to the comparative analysis of cellular processes with different conditions, CAPITAL will contribute to the elucidation of time-course regulatory systems.

## Acknowledgements

This work was supported by Japan Society for the Promotion of Science KAKENHI [18K11526 to Y. Kato; 19K20399 to T. Mori]. This study was partly achieved through the use of large-scale computer systems at the Cybermedia Center, Osaka University, and the NIG supercomputer at ROIS National Institute of Genetics.

## Code availability

CAPITAL Python codes are available at https://github.com/ykat0/capital.

